# Antibody-dependent Intracellular neutralisation by TRIM21 is potentiated by HOIP and linear ubiquitin chains

**DOI:** 10.1101/2024.04.02.587719

**Authors:** Christopher Green, Konstanze F. Winklhofer, Adam J. Fletcher, Leo C. James, John Skidmore, James Duce, William A. McEwan

**Affiliations:** UK Dementia Research Institute, University of Cambridge, Cambridge, UK; Department Molecular Cell Biology, Institute of Biochemistry and Pathobiochemistry, Ruhr University Bochum, Germany; MRC-University of Glasgow Centre for Virus Research, University of Glasgow, Glasgow, UK; MRC Laboratory of Molecular Biology, Cambridge, UK; The ALBORADA Drug Discovery Institute, University of Cambridge, Cambridge, UK

## Abstract

Antibody-dependent intracellular neutralisation (ADIN) promotes the rapid proteasomal degradation of viruses and other large substrates in the cytosol. It is dependent on detection of intracellular virus-bound antibodies by the Fc receptor and E3 ligase TRIM21, followed by disassembly of the virus by the unfoldase VCP/p97. It is not known how VCP is recruited to TRIM21. We performed a limited siRNA knock-down screen of known VCP adaptors to determine their involvement in ADIN. Knock-down of HOIP, the only ubiquitin E3 ligase capable of generating linear ubiquitin chains, resulted in impaired virus neutralisation. HOIPIN-8, a HOIP inhibitor, showed concentration-dependent reduction in virus neutralisation. We found that the activity of HOIP in ADIN is dependent on its ubiquitin binding domains and its PUB domain which recruits VCP. Knock-down of OTULIN, the only deubiquitinating enzyme that exclusively cleaves linear ubiquitin chains, potentiated neutralisation of the virus. Our results expand the role of HOIP and linear ubiquitin chains in proteostasis.

## Introduction

Neutralisation is the principal correlate of immunological protection against viral infection and is defined as the reduction of viral infectivity through the binding of antibodies to the virion. Whilst many neutralising antibodies function by preventing virus entry to the cell, the existence of a cytosolic neutralisation mechanism has recently been revealed which provides a critical last line of defence against infection. Termed antibody-dependent intracellular neutralisation (ADIN), this process relies on antibodies that bind to viral antigens that are not essential to entry. These antibodies can be brought into the cytosol in complex with the virus or subviral particle (Foss *et al*., 2019). Subsequent detection of intracellular antibodies by the cytosolic antibody receptor TRIM21 leads to rapid activation of a degradation response that neutralises infection (Mallery *et al*., 2010; McEwan *et al*., 2013).

TRIM21 is expressed in the cytosol of most cell types and has sub-nanomolar affinity for IgG. TRIM21 also possesses E3 ligase activity owing to an N-terminal RING domain. Upon binding virus:antibody complexes, the RING domain becomes active, and promotes autoubiquitination of TRIM21 with the synthesis of K63-linked ubiquitin chains. Antibody and antibody-bound targets may also become ubiquitinated by TRIM21, consistent with an urgent and acute danger signal that intracellular antibody-opsonised particles represent (Kiss *et al*., 2022; Mevissen *et al*., 2023). Following TRIM21-mediated ubiquitination, degradation of the whole antibody:virus:TRIM21 complex occurs. Infection with high titres of virus:antibody results in attendant activation of NF-κB signalling, and the secretion of proinflammatory cytokines occurs (Mallery *et al*., 2010; McEwan *et al*., 2013; Fletcher *et al*., 2015).

While initially discovered as a mechanism against non-enveloped viruses such as adenovirus, it has been demonstrated that this pathway can also degrade pathological protein aggregates such as tau fibrils during immunotherapy against neurodegenerative disorders (McEwan *et al*., 2017; Mukadam *et al*., 2023). This broad substrate specificity has been exploited in the development of an experimental technique called Trim-Away to deplete cytosolically accessible proteins as a knock-down tool (Clift *et al*., 2017). The TRIM21 pathway therefore represents an opportunity not only to degrade biomedically important targets, but also as a system to uncover how the mammalian cytosol degrades challenging and diverse proteaceous substrates. An important distinction of this pathway is how rapidly large substrates are degraded, which may have arisen due to the limited time available before irreversible infection occurs.

VCP/p97 is an abundant homohexameric unfoldase. It requires ATP to mechanically pull substrates through its central channel, prior to degradation at the proteasome (Olszewski *et al*., 2019). The 26S proteasome poorly unfolds and degrades tightly folded substrates that lack an unstructured initiation region, even if the substrates are ubiquitinated. The activity of VCP thus assists the proteasome by pre-processing substrates, permitting proteasomal access to polypeptides and potentiating degradation. VCP has a wide array of functions across the cell including roles in endoplasmic reticulum associated degradation (ERAD), mitophagy, genomic stability and ribosome associated degradation (Meyer *et al*., 2012). Its activity is regulated by adaptors that bind at the VCP N- and C-termini and specify its targets and, accordingly, its cellular role (Meyer and Weihl, 2014). It has been shown that VCP is required for the neutralisation of adenovirus during ADIN (Hauler *et al*., 2012; Mevissen *et al*., 2023), however the molecular details of this activity remain uncertain. We here sought to identify the adaptor for VCP during ADIN and found an important role for the linear ubiquitin chain-specific E3 enzyme HOIL-1 interacting protein (HOIP).

HOIP is the only ubiquitin E3 ligase in the human genome capable of generating linear (M1) ubiquitin chains (Kirisako *et al*., 2006). It is part of the heterotrimeric complex, LUBAC (linear ubiquitin chain assembly complex), alongside HOIL-1 and SHARPIN (Gerlach *et al*., 2011; Ikeda *et al*., 2011; Tokunaga *et al*., 2011). M1 chains form when the C-terminal glycine of the donor ubiquitin is covalently attached to the N-terminus of the acceptor ubiquitin (Dittmar and Winklhofer, 2020; Fiil and Gyrd-Hansen, 2021; Jahan *et al*., 2021). Unlike other ubiquitin linkages, M1 linkages are eupeptide bonds via the methionine amino group which can be cleaved by two deubiquitinating enzymes (DUBs), CYLD which is bi-specific for M1 and K63 chains, and OTULIN which solely cleaves M1 chains (Keusekotten *et al*., 2013; Elliott *et al*., 2021).

Structurally, HOIP consists of an N terminal PUB domain which promotes interaction with SPATA2, OTULIN and VCP (Elliott *et al*., 2014; Schaeffer *et al*., 2014; Kupka *et al*., 2016). This is followed by NZF domains which bind to ubiquitin chains and have high affinity for K63 ubiquitin chains (Emmerich *et al*., 2013) and K48 chains (Tobias L. Haas *et al*., 2009; Ikeda, *et al*., 2011). Finally, at the C terminus is the RBR E3 ligase domain followed by a linear determination domain (LDD) which is essential for the formation of M1 ubiquitin chains (Smit *et al*., 2012; Stieglitz *et al*., 2013).

The role for M1 chains was initially thought to be isolated to immune signalling. It has been demonstrated that interaction between linear ubiquitin chains and NEMO (NF-κB essential modifier)/IKKɣ via its UBAN domain results in upregulation of NF-κB signalling (Rahighi *et al*., 2009; Tobias L Haas *et al*., 2009; Fujita *et al*., 2014). NEMO can also be modified with M1 chains promoting its oligomerisation and the trans-autophosphorylation of IKK⍺ and IKKβ (Tokunaga *et al*., 2009; Ikeda *et al*., 2011; Rahighi *et al*., 2022). Recently the roles of HOIP and M1 chains have been expanded to include proteostasis. HOIP is recruited to cytoplasmic protein aggregates through binding to the PIM domain of VCP/p97 by its PUB domain, and *vice versa*, VCP/p97 abundance at aggregates is increased by HOIP (Well *et al*., 2019). As a result, the proteotoxicity of various protein aggregates, such as huntingtin or synuclein aggregates is reduced (Well *et al*., 2019; Furthmann *et al*., 2023).

Here we sought to identify and validate the adaptor responsible for recruiting VCP to TRIM21 and thus providing the ADIN pathway with the capacity to rapidly degrade challenging substrates. Our results implicate HOIP and M1 chains as factors that potentiate virus neutralisation during ADIN.

## Results

TRIM21 mediates a selective degradation response against diverse antibody-bound substates with the involvement of VCP. To better understand this potent degradation pathway, we sought to identify the adaptor(s) that promote VCP recruitment to activated TRIM21. We used a well-established adenovirus neutralisation assay as a platform for a targeted RNAi screen using siRNA sequences specific for 31 genes with established roles as VCP adaptors. During this assay, replication-deficient human adenovirus serotype 5 encoding eGFP under a CMV promoter (AdV) is pre-incubated with either vehicle (PBS) or with anti-hexon antibody 9C12. 9C12 is a non-entry blocking monoclonal antibody which neutralises AdV infection in a TRIM21-dependent manner (McEwan *et al*., 2012). We first sought to confirm the role of VCP in ADIN in HEK293 wildtype (WT) cells, a widely used human cell line highly amenable to RNAi. HEK293 WT cells were treated with 10 nM siRNA for 48 hours followed by infection with AdV pre-incubated with 9C12 for one hour at room temperature (or PBS as an infection control). 24 hours after infection, cells were collected and GFP expression (proxy for infection) was quantified by flow cytometry. Pre-incubation of AdV with 9C12 resulted in an antibody concentration dependent reduction in infection and corresponding neutralisation of virus infection (*Fig 1A & B*). Neutralisation was substantially reduced when either TRIM21 or VCP were depleted by siRNA compared to a non-targeting control (NTC) siRNA (*Fig 1C & D*), confirming the cell type as suitable for a VCP adaptor screen. Protein knock-down efficiency was confirmed by immunoblot (*Fig 1E*).

**Figure 1:**
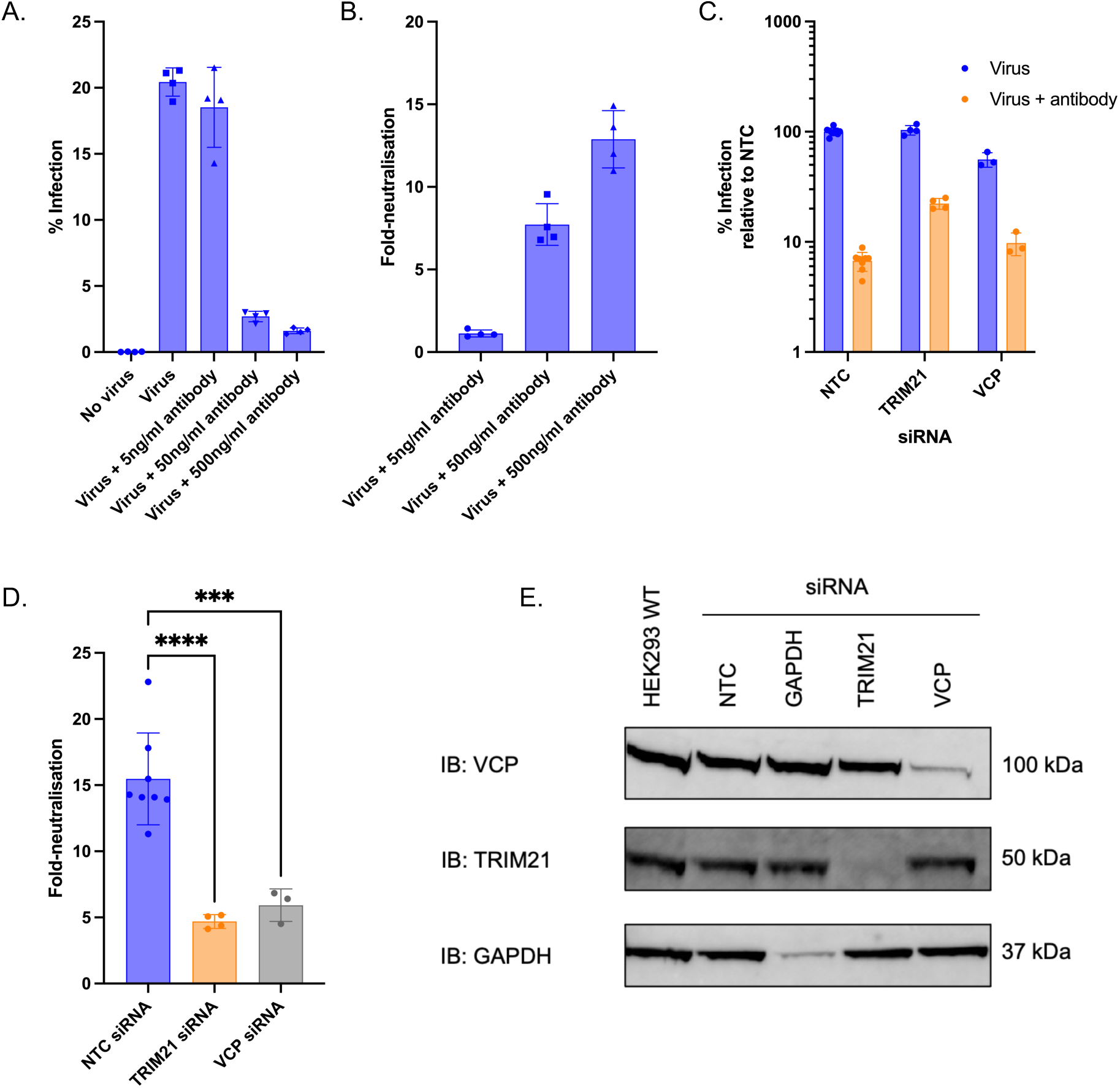
Infection levels in HEK293 cells treated with a titration of antibody (9C12). Individual data points indicated +/- S.D. (A). Same data as A but data converted to fold-neutralisation by dividing each of the antibody conditions by the mean of the virus only condition. Individual data points indicated +/- S.D. (B). Infection levels of HEK293 with or without TRIM21 or VCP siRNA treatment for 48 hours, followed by infection with virus +/- antibody (C). Same data as in C but converted to fold-neutralisation, +/- S.D. (D). Immunoblot of HEK293 WT cells treated with 10 nM of indicated siRNA for 72 hours (E). Statistical statement: One-way ANOVA with multiple comparisons to NTC *P <0.05 **P < 0.01; ***P < 0.001; ****P < 0.0001 (D).

We next treated cells with pooled siRNA targeting 31 known VCP adaptors and performed AdV neutralisation assays. The majority of proteins yielded no statistically significant effect on neutralisation. Treatment with siRNA targeted to four genes besides TRIM21 yielded a statistically significant reduction (P<0.05) in neutralisation: UBXN1, HOIP, UBXD3 and UBXD8. A further two demonstrated an increase in neutralisation compared to control conditions, VIMP and UBXD1. Of the siRNA treatments that reduced neutralisation, two provided a phenotype comparable in magnitude to TRIM21 siRNA: UBXN1 and HOIP *(Fig 2A*) and were taken forward for validation. Cells treated with HOIP siRNA showed reduced HOIP expression and unchanged levels of UBXN1 expression (*Fig 2D*). Cells treated with UBXN1 siRNA likewise showed reduced UBXN1 expression, but also unexpectedly demonstrated reduced HOIP expression. This suggests that the decrease in neutralisation in UBXN1 siRNA treated cells is likely due to a reduction in HOIP expression. Thus, UBXN1 may have a role in regulation of HOIP, but is less likely to be directly involved in ADIN. As such, the role of HOIP and M1 ubiquitin chains in ADIN became the focus of this study. Validated knockdown of HOIP resulted in increased infection by AdV in complex with 9C12 while having little effect on naked AdV (*Fig 2B*), similar to the effect of TRIM21 knockdown, resulting in a significant impairment of virus neutralisation (*Fig 2C*). This suggests that HOIP, the linear ubiquitin E3 ligase and catalytic component of the LUBAC complex, has a previously unknown role in ADIN.

**Figure 2:**
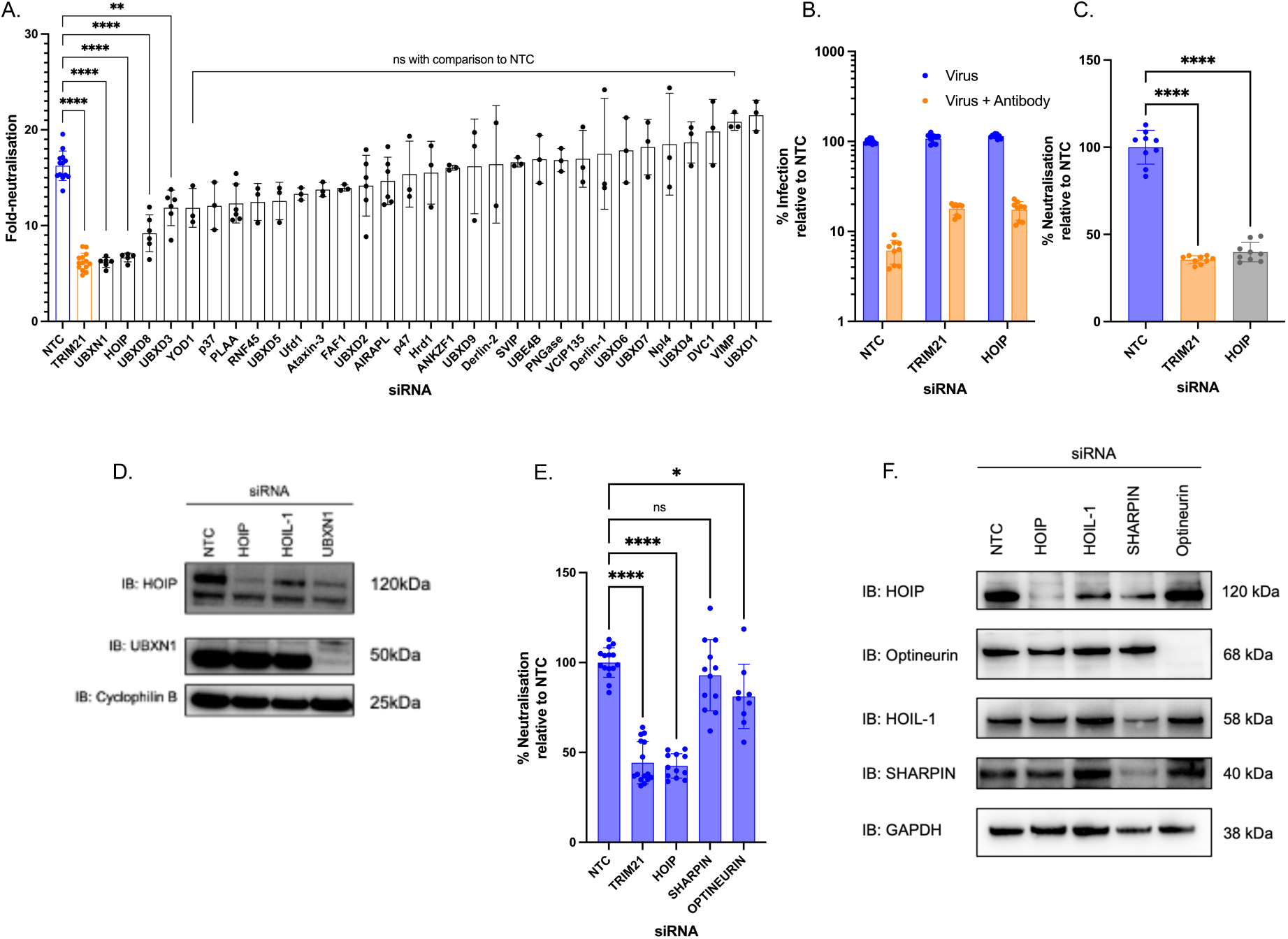
Fold-neutralisation of adenovirus infection by 9C12 under conditions of siRNA treatment. siRNA targets are listed as protein names or non-targeting control (NTC), individual data points plotted, bars indicate mean +/- S.D. (A). Infection of HEK293 WT cells treated with indicated siRNA for 48 hours followed by infection with virus +/- antibody for 24 hours, bars indicate mean +/- S.D. (B). Fold neutralisation data from 1B, (C). Immunoblot of HEK293 cells treated with 10 nM indicated siRNA for 72 hours (D). Fold-neutralisation of infection in HEK293 WT cells treated with indicated siRNA for 48 hours followed by virus +/- antibody for 24 hours, bars indicate mean +/- S.D. (E). Immunoblot of HEK293 cells treated with 10 nM indicated siRNA for 72 hours (F). Statistical statement: One-way ANOVA with multiple comparisons to NTC *P <0.05 **P < 0.01; ***P < 0.001; ****P < 0.0001 (A, C & E).

LUBAC contains three proteins, HOIP, HOIL-1 and SHARPIN. We therefore sought to determine the role of the other LUBAC components in ADIN. SHARPIN knockdown had no significant effect on ADIN compared to NTC suggesting it is dispensable for ADIN (*Fig 2E*). We also tested Optineurin, an autophagy adaptor that targets substrates modified with M1 ubiquitin chains for degradation by autophagy (Wild *et al*., 2011; Wong and Holzbaur, 2014). We observed a mild but significant reduction in neutralisation upon treatment with Optineurin siRNA. This indicates that while the proteasome has been established as the main route for degradation in this pathway, there may remain a further role for autophagy (*Fig 2E)*. Immunoblots of HEK cells treated with siRNA for 72 hours showed efficient knock-down of most protein targets (*Fig 2F*). Notably HOIL-1 siRNA appeared to fail to reduce expression of HOIL-1 and therefore was not analysed for its role in neutralisation.

HOIP is a large protein comprising multiple domains with defined functions and binding capabilities. We next sought to determine which HOIP domains are required for ADIN. To do this, we used complementation of HOIP deficient cells. We treated HEK293 cells with CRISPR gRNA constructs targeting *RNF31*, the gene encoding HOIP. To validate successful targeting of this gene, we treated cells with TNF-⍺ and measured the phosphorylation of IκB⍺. It has been established that LUBAC and linear ubiquitination are required to transduce the signal from activation of the TNF receptor TNFR1 to NF-κB activation (Tobias L Haas *et al*., 2009; Tokunaga *et al*., 2009; Peltzer *et al*., 2014). We observed a strong increase in p-IκB⍺ following TNF⍺ treatment in HEK293 WT cells, however in the HEK293 HOIP KO line, this phosphorylation was delayed and attenuated (*Fig 3A)*, consistent with the function of LUBAC being decreased. Additionally, an SDS-PAGE band detected by an anti-HOIP antibody at 120 kDa was severely diminished in these cells, consistent with expression of full-length HOIP being attenuated.

**Figure 3:**
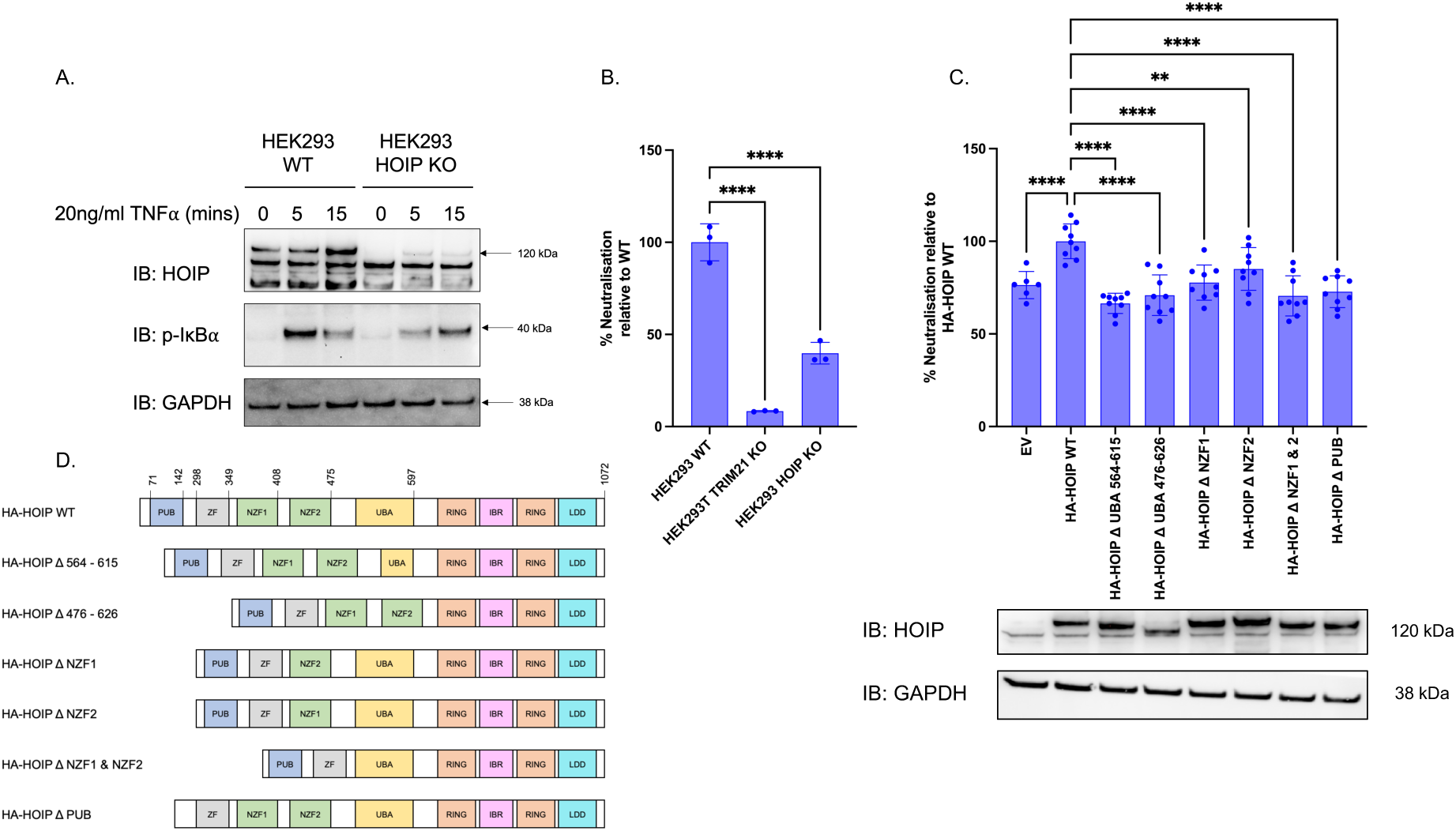
HEK293 WT and HEK293 HOIP knock-out cells were treated with 20 ng/ml TNF-⍺ for the indicated timepoints, cells were lysed and analysed by immunoblot (A). HEK293 WT, HEK293T TRIM21 knock-out and HEK293 HOIP knock-out cells were infected with either virus or antibody pre-incubated virus for 24 hours and fold-neutralisation calculated. Data is % relative to HEK293 WT, bars indicate mean +/- S.D. (B). HEK293 HOIP depleted cells were transfected with indicated constructs for 72 hours, followed by infection with either virus or antibody pre-incubated virus for 24 hours. Fold-neutralisation was calculated, data relative to HA-HOIP WT, bars indicate mean +/- S.D. Below – corresponding immunoblots for the 72 hour transfection timepoint (C). HOIP constructs with deletions indicated, domains not to scale (D). Statistical statement: One-way ANOVA with multiple comparisons to NTC *P <0.05 **P < 0.01; ***P < 0.001; ****P < 0.0001 (B & C).

Next, we tested the HOIP cells in the adenovirus neutralisation assay and, consistent with data from siRNA, observed significantly lower levels of virus neutralisation.

When compared against HEK293T TRIM21 KO cells, HOIP KO cells showed a lower degree of neutralisation impairment. This suggests that HOIP is not essential for ADIN, but acts to increase the efficiency of TRIM21 mediated neutralisation (*Fig 3B*).

To determine which HOIP domains are necessary in ADIN, we complemented the HEK293 HOIP KO cell line with a panel of plasmids (Well *et al*., 2019) encoding HOIP constructs with the indicated domain deletions (*Fig 3D*). Full-length HOIP constructs significantly increased virus neutralisation compared to empty vector, confirming that restoration of HOIP expression was able to exert partial rescue of ADIN. The UBA domain of HOIP interacts with the UBL domain of HOIL-1 and SHARPIN and these interactions are required to release autoinhibition of HOIP (Smit *et al*., 2012; Stieglitz *et al*., 2012; Liu *et al*., 2017). Total deletion of UBA (HOIP-Δ1476-626) and partial deletion of the region (HOIP-Δ1564-615) are both expected to ablate the interaction with HOIL-1 and SHARPIN. We observed that both of these constructs were unable to rescue the ADIN phenotype, consistent with interaction between HOIL-1 or SHARPIN and HOIP being essential to the role of HOIP in ADIN (*Fig 3C*).

LUBAC is recruited to its targets via interactions between its NZF domains and ubiquitin chains. To determine whether this activity was required for effective ADIN, we used constructs with one or both NZF domains deleted. The HOIP-Δ1NZF1, HOIP-Δ1NZF2 and HOIP-Δ1NZF1 & NZF2 domain deletion constructs failed to rescue neutralisation, confirming the importance of the NZF domains in ADIN.

HOIP recruits VCP to its targets via its N-terminal PUB domain. Consistent with this activity being essential for HOIP activity in ADIN, we found that HOIP-ι1PUB was unable to rescue neutralisation. This observation is consistent with the status of HOIP as a VCP adaptor in this pathway. For all deletion constructs, immunoblot confirmed that expression was reasonably consistent (*Fig 3C*).

The results thus far demonstrate that HOIP is involved in the process of ADIN. We hypothesised that its established catalytic activity, namely synthesis of M1-linked ubiquitin chains, may therefore be essential for full levels of TRIM21-mediated neutralisation. To test the role of M1 chains in ADIN, we used HOIPIN-8, a compound which specifically inhibits linear chain generation by binding to residues in the catalytic RBR and LDD domains of HOIP (Oikawa *et al*., 2020). Through this binding, HOIPIN-8 acts as an inhibitor of M1 chain formation. We saw concentration-dependent inhibition of virus neutralisation in cells treated with HOIPIN-8 consistent with M1 chain formation being required for effective ADIN (*Fig 4D*). The observed IC_50_ was 549 nM, consistent with the published cellular IC_50_ value for this drug of 420 nM (Oikawa *et al*., 2020). Our results thus implicate the catalytic activity of HOIP as an important contributor to ADIN.

**Figure 4:**
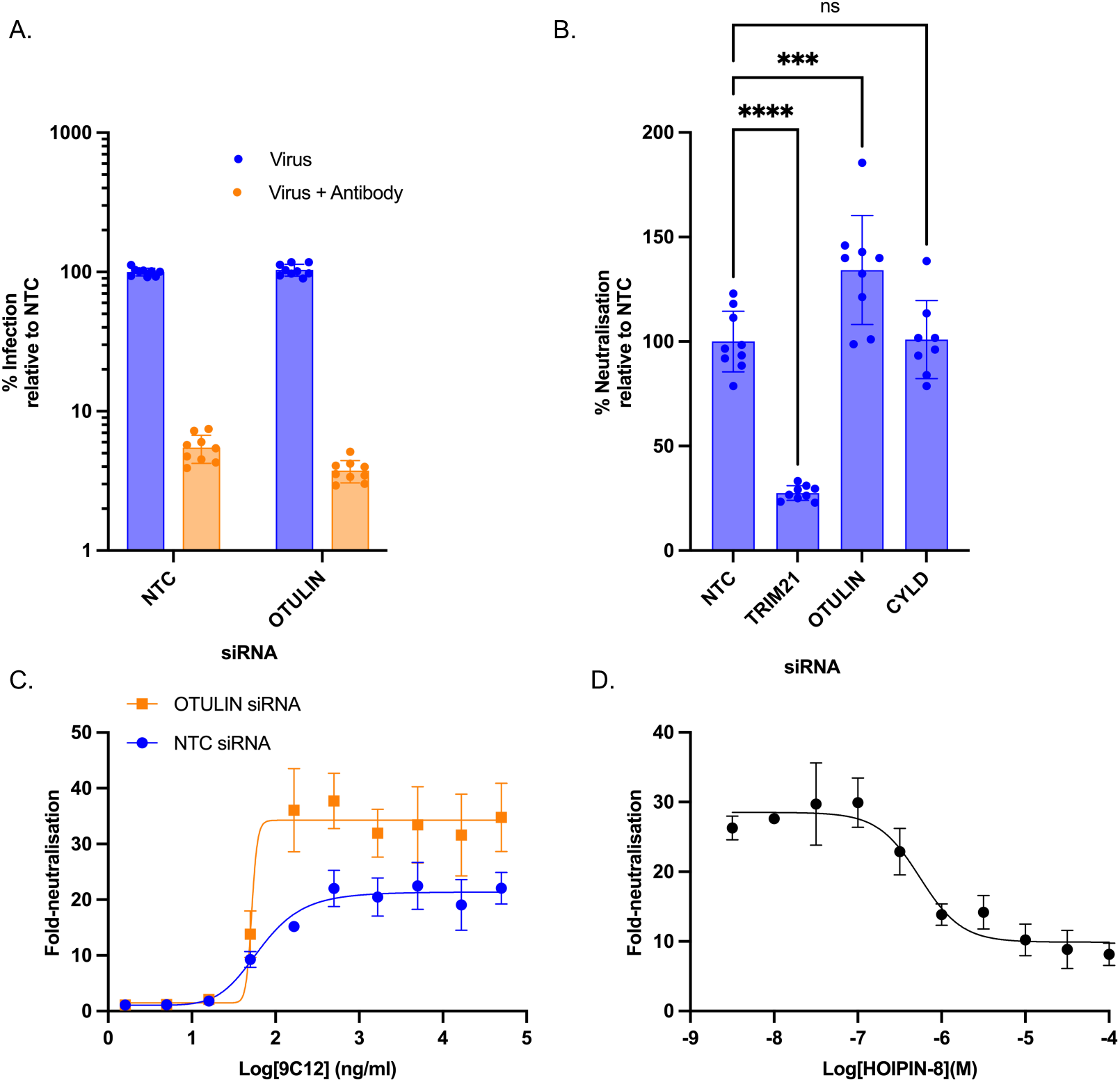
Infection of HEK293 WT cells treated with indicated siRNA for 48 hours followed by infection with virus +/- antibody for 24 hours, bars indicate mean +/- S.D. (A). Fold neutralisation data from 4A, data relative to NTC, bars indicate mean +/- S.D. (B). HEK293 WT cells were treated with indicated siRNA at 10 nM for 48 hours then infected with virus pre-incubated with a half-log antibody dilution. Note the highest concentration is 50 µg/ml. Fold-neutralisation calculated as before (C). HEK293 WT cells were pre-incubated with the indicated HOIPIN-8 concentrations for 1 hour, followed by infection with either virus only, or virus pre-incubated with antibody (D). Statistical statement: One-way ANOVA with multiple comparisons to NTC *P <0.05 **P < 0.01; ***P < 0.001; ****P < 0.0001 (B).

Our results are consistent with a model wherein LUBAC stimulates M1 chain synthesis in order to stimulate efficient degradation of antibody:virus complexes. A prediction of this model is that depletion of M1 deubiquitinating enzymes (DUBs) will enhance this activity and permit greater levels of ADIN. M1 chains are degraded by the DUBs CYLD, which can degrade M1 or K63 chains, and OTULIN, which has exclusive activity for M1 chains. We performed the adenovirus neutralisation assay in HEK293 WT cells treated with siRNA targeting these DUBs. We found that reduction of CYLD expression had no significant effect on neutralisation compared to the non-targeting control siRNA. However, we found that by reducing expression of OTULIN we potentiated the ability of the cell to neutralise virus infection (*Fig 4B*). This effect was driven by reduced infection in the antibody treated condition (*Fig 4A*). Of note, OTULIN depletion did not improve the EC_50_ of the antibody-neutralisation, with observed 9C12 EC_50_ being 68 ng/ml for NTC and 52 ng/ml for OTULIN knock-down (*Fig 4C*). Rather, its effects were manifest at high concentrations of antibody, in the so-called ‘persistent fraction’ of virus infection, the level of virus infection that remains at high concentrations of antibody. Previous findings have shown the persistent fraction to be dependent on cellular TRIM21 concentration levels (McEwan *et al*., 2012). This is the first demonstration, to our knowledge, that the persistent fraction can be lowered by altering anything other than TRIM21 itself. The results are consistent with LUBAC being a co-factor for ADIN and OTULIN acting as its negative regulator.

## Discussion

Antibody-dependent intracellular neutralisation (ADIN) is a pathway that promotes the proteasomal degradation of antibody-bound virus in the intracellular space. It is dependent on binding of antibodies by the cytosolic Fc receptor and E3 ubiquitin ligase TRIM21. Viruses are large and potentially challenging substrates for cellular degradative pathways. VCP is recruited to the TRIM21:antibody:virus complex likely to assist unfolding and separation of the complex to promote its degradation. We here used a limited siRNA screen of VCP adaptors and identify HOIP as an important cofactor to ADIN. HOIP makes a suitable candidate as it is recruited by interactions between its NZF domains and K63 ubiquitin chains, the enzymatic product of TRIM21. Like TRIM21, HOIP is broadly expressed in most cell types. Its genetic targeting using CRISPR/Cas9 rendered cells poorly able to execute the ADIN pathway, resulting in increased vulnerability to infection by antibody-coated adenovirus. Our results reveal a new and unexpected role for HOIP in selective degradation pathways.

Analysis of the HOIP domains that contribute to ADIN demonstrated essential functions for the PUB domain (VCP recruitment), NZF1 domain (ubiquitin binding) and UBA domain (HOIL-1/SHARPIN binding). Furthermore, E3 ligase activity of HOIP was essential as inhibition of HOIP RBR domain using HOIPIN-8 led to a dose-dependent reduction in ADIN activity. We therefore propose a model wherein TRIM21 synthesises K63-linked ubiquitin chains following engagement of antibody-bound virus particles in the cytosol and LUBAC is recruited to these sites of ADIN owing to its affinity for K63 chains and contributes M1-linked ubiquitin. This M1 modification enhances ADIN, as ADIN can be both reduced by inhibition of M1 chain formation (with HOIPIN-8) and enhanced by stabilization of M1 chains (OTULIN depletion) respectively.

It remains uncertain why the E3 ligase activity of HOIP increases the efficiency of ADIN given that an adaptor activity of HOIP could in principle occur without it. Future research should be guided by findings that HOIL-1 can be recruited to M1 ubiquitin chains (Tobias L. Haas *et al*., 2009; Gomez-Diaz *et al*., 2021) thereby potentially initiating a feed-forward amplification of LUBAC recruitment and its VCP adaptor function. A further area of research could be the contribution of HOIP and M1 chains to the secondary immune signalling observed during ADIN. This is particularly pertinent due to the well-established role of HOIP and M1 chains in immune signalling.

Passive antibody therapy against misfolded protein tau is, like certain viral infections, susceptible to neutralisation by TRIM21 (Mukadam *et al*., 2023). This raises the question of whether linear ubiquitination would also be involved in the degradation of other substrates, in particular misfolded tau. Interestingly, it has been shown that an interaction between LUBAC and huntingtin aggregates promotes VCP recruitment and is a protective event against the accumulation of aggregates (Well *et al*., 2019). Moreover, M1 ubiquitin chains are enriched at tau, ⍺-synuclein and TDP-43 aggregates in patients’ brain (Nakayama *et al*., 2019, 2020; Furthmann *et al*., 2023). Thus, similar pathways may be employed during homeostatic control of protein aggregation and acute innate immune responses to antibody-opsonised intracellular particles. Understanding these pathways and their mechanistic differences may therefore reveal new and unexpected means by which innate immunity can be stimulated to selectively degrade protein aggregates.

Some limitations of our research remain. Protein-protein interaction studies of ADIN are particularly difficult to perform. Indeed currently, co-immunoprecipitation of TRIM21 requires over-expression of His-tagged TRIM21 and N-terminal HA-tagged ubiquitin (Fletcher *et al*., 2015). While this successfully demonstrated the presence of K48 and K63 chains on TRIM21, the N-terminal tag may interfere with M1-linked ubiquitination. As a result of this limitation, we have not demonstrated that TRIM21 is modified with M1 chains, nor that VCP is directly recruited by HOIP. A recent publication has used *Xenopus* egg extract to study ADIN and has detected chain specific ubiquitination of TRIM21 and demonstrated VCP-dependence (Mevissen *et al*., 2023). Such *in vitro* reconstitution systems could be adapted to study the role of HOIP and M1 chains, particularly if a challenging, complex substrate is the target for degradation.

In summary, we have identified HOIP as a cofactor for intracellular virus neutralisation via TRIM21. Ablation of HOIP resulted in impaired viral neutralisation, as did pharmacological inhibition of M1 chain formation. Conversely, inhibition of M1 chain hydrolysis exerted an opposite effect on overall levels of neutralisation. Our results reveal an unexpected role for HOIP and M1 chains in acute antiviral protein degradation responses.

## Materials and Methods

### Cell Culture

HEK293 WT, HEK293T TRIM21 knock-out and HEK293 HOIP KO cells were maintained at 37°C in complete DMEM (ThermoFisher 10569010) supplemented with 10% FBS and pen/strep.

### HEK293 HOIP CRISPR Knock-out

Guide RNA (Hs.Cas9.RNF31.1.AS – sequence: CAGGAGCAATCTCTCTCAAT targeting exon 1), Alt-R CRISPR-Cas9 tracrRNA (1073190) and Alt-R S.p. Cas9 Nuclease V3 (1081058) were purchased from IDT. gRNA complex was prepared by mixing 5µl 100µM guide with 5µl 100µM tracrRNA and heated at 95°C for 5 minutes. Recombinant Cas9 was diluted to 1µM in OptiMEM. 1.5µl gRNA complex and 1.5µl diluted Cas9 was mixed with 22µl OptiMEM and incubated at room temperature for 5 minutes to form the RNP complex. 1.2µl Lipofectamine RNAiMAX and 23.8µl OptiMEM was added to the RNP complex followed by a 20 minute incubation at room temperature. HEK293 WT cells were diluted to 400,000 cells/ml in DMEM with 10% FBS (no antibiotics). 50µl of transfection mix was added to a well of a 96-well plate and 100µl cells were added on top. Cells incubated at 37°C for 48 hours. Knock-out was analysed by immunoblot. A single cell clonal expansion was performed to generate a monoclonal cell line.

### Adenovirus Neutralisation Assay

HEK293 cells were diluted to 200,000 cells/ml in complete DMEM (no antibiotics). 500µl of cells was added to each well of a 24 well plate (Greiner 662160) and incubated at 37°C overnight. Replication-deficient adenovirus (human, serotype 5) encoding GFP (hAdV5 GFP) under a CMV promoter (ViraQuest Inc - VQAd CMV eGFP-2.6del 080420) was diluted to 1x10^8.52^/ml in PBS and incubated with either 500ng/ml 9C12 or PBS at room temperature for 1 hour. 20µl of either virus only or virus and antibody was added to each well, roughly 333,333 viral units/well. 20µl of either virus only or virus and antibody was added to each well. The plate was incubated at 37°C for 24 hours. The next day media was removed and cells washed with 250µl PBS. PBS was removed and 150µl 0.25% trypsin was added to each well and the plate briefly incubated at 37°C. Once the cells had detached 150µl DMEM (with 10% FBS) was added to each well and the 300µl volume from each well was transferred to a clear U-bottom 96-well plate. The plate was centrifuged at 300xG for 10 minutes at 4°C. Media was removed and cells resuspended in 150µl 4% PFA v/v in PBS. Plates were sealed and stored in the dark at 4°C until quantifying GFP expression by flow cytometry on the Beckman CytoFLEX.

#### For siRNA knock-downs

The siRNA was diluted to 50nM in OptiMEM with 1% Lipofectamine RNAiMAX. The mix was incubated at room temperature for 15 minutes. 125µl of siRNA mix was added to each well of a 24-well plate. HEK293 WT cells were diluted to 150,000 cells/ml in DMEM (+10% FBS, no antibiotics). 500µl of cells was added to each well. The plate was incubated at 37°C for 48 hours. Following this virus and antibody treatment was performed as above for 24 hours.

#### For HOIP construct transfection

0.5µg DNA was mixed with 400µl OptiMEM followed by addition of 1µl P3000 reagent. 45µl of Lipofectamine 3000 was added to 4ml OptiMEM. Diluted DNA and Lipofectamine were mixed 1:1 (400µl + 400µl) and incubated at room temperature for 15 minutes. HEK293 HOIP knock-out cells were diluted to 150,000 cells/ml in antibiotic-free DMEM (with 10% FBS). 125µl transfection mix was added to each well of a 24-well plate followed by 500µl cells. The plate was incubated at 37°C for 72 hours followed by the virus and antibody treatment as above for 24 hours.

### Immunoblot

Cells were washed with ice-cold PBS then lysed in lysis buffer (20mM Tris HCl (pH8,0), 137mM NaCl, 1% NP-40, 2mM EDTA, supplemented with protease and phosphatase inhibitors) on ice for 20 minutes with regular agitation. Samples were centrifuged at 12,000 x G for 20 minutes at 4°C and the supernatant retained. Reducing laemelli reagent was added to the sample and heated at 100°C for 5 minutes. Samples were typically ran on a 4-12% Bis-tris gel at 200V for 40 minutes using MES running buffer. Proteins were transferred to a PVDF membrane using the Biorad Trans-blot Turbo. Membranes were blocked with 5% milk in TBST (0.1% tween) for 1 hour at room temperature. Primary antibodies were diluted in blocking buffer and incubated with the membrane overnight at 4°C. Membranes were washed with TBST three times, ten minutes each. Secondary antibodies were diluted in blocking buffer and incubated with the membrane at room temperature for 1 hour. Membranes were washed TBST three times, ten minutes each and imaged on the Biorad Chemidoc.

### Antibodies and Reagents

| Target | Company | Catalog Number | Use |
| --- | --- | --- | --- |
| TRIM21 | ProteinTech | 12108-1-AP | Immunoblot |
| HOIP | Abcam | ab125189 | Immunoblot |
| VCP | Genetex | GTX101089 | Immunoblot |
| UBXN1 | Genetex | GTX121127 | Immunoblot |
| GAPDH | ThermoFisher | MA5-15738 | Immunoblot |
| p-IkBα | Cell Signalling | 2859 | Immunoblot |
| Cyclophilin B | Santa Cruz | Sc-390193 | Immunoblot |
| HOIL-1 | Abcam | ab108479 | Immunoblot |
| SHARPIN | Abcam | ab236092 | Immunoblot |
| Optineurin | Abcam | ab213556 | Immunoblot |

| Secondary | Company | Catalog Number |
| --- | --- | --- |
| Goat anti-mouse IgG HRP | BioRad | 1721011 |
| Goat anti-rabbit IgG HRP | BioRad | 1706515 |
| Goat anti-mouse IgG2 alexa 488 | Thermo | A-21131 |
| Goat anti-rabbit alexa 568 | Thermo | A-11011 |
| Goat anti-mouse IgG1α alexa 647 | Thermo | A-21240 |

9C12 (mouse anti-adenovirus hexon) was produced by hybridomas and purified in-house following a published protocol (McEwan *et al*., 2012).

### Individual siRNA

Note: all siRNA was purchased from Horizon Discovery. All siRNA are ON-TARGETplus SMARTpool.

| Target | Catalog Numbers |
| --- | --- |
| VCP | L-008727-00-0005 |
| TRIM21 | L-006563-00-0005 |
| Non-targeting control | D-001810-10-05 |
| HOIP | L-021419-00-0005 |
| UBXN1 | L-008652-01-0005 |
| HOIL-1 | L-006932-00-0005 |
| SHARPIN | L-018575-02-0005 |
| Optineurin | L-016269-00-0005 |
| CYLD | L-004609-00-0005 |
| OTULIN | L-018991-02-0005 |

## Acknowledgments

CG was funded by an ARUK PhD studentship (ARUK-PhD2018-051). WM was supported by a Sir Henry Dale Fellowship jointly funded by the Wellcome Trust and the Royal Society (grant 206248/Z/17/Z) and by the Lister Institute for Preventative Medicine. Further support was provided by the UK Dementia Research Institute (award number UK DRI-2010) through UK DRI Ltd, principally funded by the UK Medical Research Council. KFW’s work is supported by the German Research Foundation (WI/2111-6, WI/2111-8, FOR 2848), Germany’s Excellence Strategy EXC 2033 390677874 – RESOLV, and the Michael J. Fox Foundation for Parkinson’s Research (Grant ID 021968).

